# Engineering Heterotypic Biomolecular Condensates with Synthetic Peptides for Controlled Spatial Organization and Liquid-like Nature

**DOI:** 10.64898/2026.08.09.743201

**Authors:** Sudhriti Roy, Deepak Sharma, Milan Kumar Hazra

## Abstract

Sequence heterogeneity is a defining feature of cellular biomolecular condensates, yet how competing interaction motifs encode their thermodynamic stability, internal organization, and dynamics remains poorly understood. Here, we systematically tune the hydrophobicity mismatch between intrinsically disordered peptide pairs to establish sequence hydrophobicity as a programmable determinant of heterotypic condensate behaviour. We show that heterotypic condensates are thermodynamically more stable than homotypic ones having same average hydrophobicity through the cooperative interplay of short-range hydrophobic and long-range electrostatic interactions. Increasing hydrophobicity mismatch drives a composition-dependent transition from homogeneous condensates to core–shell architectures accompanied by pronounced spatial and dynamical heterogeneity, whereas reducing sequence disparity restores homogeneous organization and nearly uniform dynamics. Our results establish a direct molecular link between sequence chemistry, phase stability, condensate architecture, and transport dynamics, providing predictive design principles for engineering synthetic biomolecular condensates with programmable organization and material properties.

**TOC:** 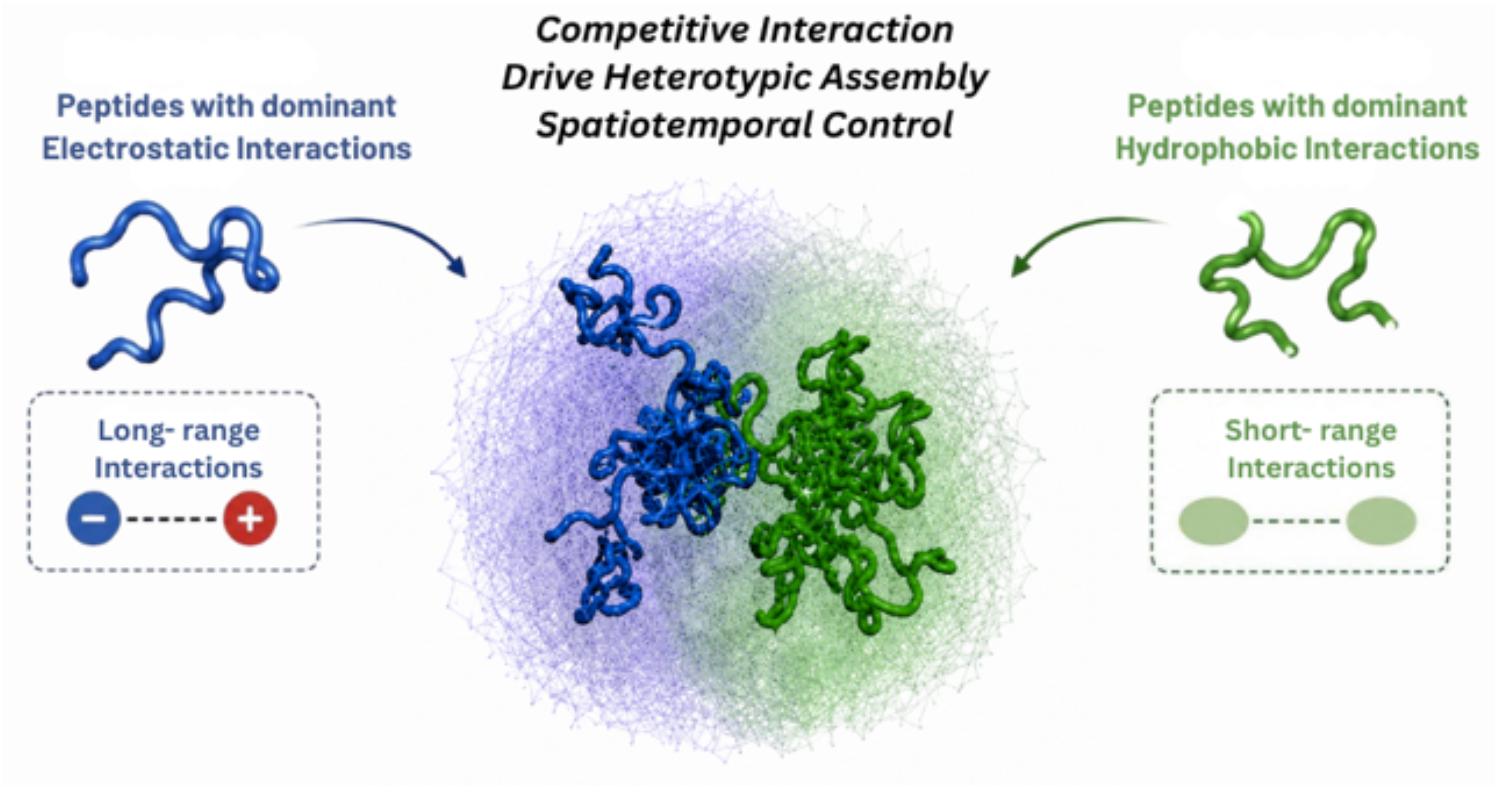

## Introduction

Biomolecular condensates formed through liquid–liquid phase separation provide a fundamental mechanism for organizing the intracellular environment without membrane boundaries^1–6^. These dynamic assemblies regulate diverse cellular processes including stress adaptation^7–11^, ribosome biogenesis^12^, chromatin organization^13^, transcriptional control^14–16^, and DNA damage repair^17^, while aberrant phase separation has been increasingly linked to neurodegenerative disorders^18–22^ and other protein aggregation diseases^22–24^. A central challenge in the field is to establish sequence-level design principles that enable rational control over condensate stability, internal organization, and material properties, thereby facilitating the engineering of synthetic condensates with programmable functions^25–28^.

Intrinsically disordered proteins are particularly prone to phase separation^29–34^ because their flexible conformations support multivalent intermolecular interactions^35–45^ arising from electrostatic^46^, hydrophobic^47,48^, π-mediated^49^, and hydrogen-bonding contacts. Although homotypic phase separation has been extensively investigated, most intracellular condensates are multicomponent assemblies^50–54^ in which heterotypic interactions between biomolecules possessing distinct sequence chemistries dictate condensate formation and function. The balance between self- and cross interactions governs molecular partitioning, phase stability, and the emergence of complex condensate architectures, yet the molecular principles underlying this competition remain incompletely understood.

An intriguing feature of many cellular condensates is the presence of internal spatial organization^5,55^. The nucleolus, stress granules, P-bodies, and numerous RNA–protein condensates exhibit multilayered or multiphase architectures that compartmentalize^51,56–60^ distinct biochemical activities within a single droplet. Similar morphologies have been reproduced in vitro using mixtures of proteins, peptides, and RNA, demonstrating that relatively simple molecular systems can spontaneously generate complex mesoscale organization. Formation of such multiphase condensates, however, requires overcoming the energetic penalty associated with creating internal interfaces, implying that favourable sequence-dependent interactions must compensate for the increased interfacial free energy. Identifying the interaction motifs that stabilize these architectures therefore represents a fundamental thermodynamic problem.

Recent studies suggest that the competition between long-range electrostatic attractions and short-range hydrophobic interactions plays a dominant role in determining condensate organization^42^. Nevertheless, naturally occurring condensates contain proteins spanning a broad spectrum of hydrophobicity and charge distributions, making it difficult to isolate how differences in sequence chemistry influence phase behaviour. Consequently, a predictive understanding of how compositional heterogeneity regulates condensate stability, compartmentalization, and molecular dynamics remains lacking.

Here, we address this question by systematically engineering binary peptide mixtures with controlled hydrophobicity mismatch while independently varying their mixing composition. Using coarse-grained molecular simulations, we demonstrate that heterotypic condensates can exhibit greater thermodynamic stability than homotypic condensates with the same average hydrophobicity, with stabilization increasing as the hydrophobicity mismatch becomes larger. More importantly, the competition between short-range hydrophobic and long-range electrostatic interactions drives composition-dependent spontaneous compartmentalization. Hydrophobic peptide enrichment produces condensates with slowly relaxing hydrophobic cores surrounded by electrostatically enriched dynamic interfacial layers, whereas electrostatic enrichment reverses this organization to generate charge-dominated cores encapsulated by hydrophobic surface layers. As the hydrophobicity mismatch decreases, spatial segregation progressively disappears, yielding homogeneous condensates. Together, these results establish hydrophobicity mismatch and compositional control as sequence-encoded design parameters that govern the thermodynamic stability, internal architecture, and dynamic heterogeneity of heterotypic biomolecular condensates, providing general principles for understanding naturally occurring membrane-less organelles and for engineering programmable synthetic condensates.

To uncover how sequence-encoded hydrophobic heterogeneity governs biomolecular condensates, we designed three pairs of 40-residue intrinsically disordered peptides with progressively increasing hydrophobicity differences (Δϕ=0.1–0.5) as order parameter, while maintaining identical average hydrophobicity and overall charge neutrality. Heterotypic condensates spanning a broad range of mixing fractions were compared with homotypic condensates of equivalent composition to isolate the effects of sequence asymmetry. Equilibrium phase behaviour was investigated using a residue-level coarse-grained model^61^ that captures electrostatic and hydrophobic interactions. This framework enables direct quantification of how differential hydrophobicity reshapes condensate morphology, thermodynamic stability, and molecular mobility, establishing sequence hydrophobicity as a key molecular determinant of condensate organization and material properties.

### Simulation Details

Langevin dynamics simulations have been carried out for various mixing fractions of peptides (*X*_*High*−*ϕ*_, fraction of high-ϕ peptides in designed systems) with an embedded implicit solvent model and salt effect to screen electrostatic interactions among charged residues through Debye-Hückel potential^62^. 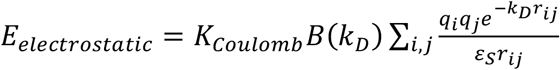 where *q*_*i*_ and *q*_*j*_ denote the charge of the *i*^th^ and j^th^ bead, *r*_*ij*_ denotes the inter-bead distance, *ε*_*S*_ is solvent dielectric constant, and *K*_*Coulomb*_ = 4*πε*_0_ =332 kcal/mol. *B*(*k*_*D*_) is a function of solvent salt concentration and the radius (*a*) of ions produced by the dissociation of the salt, and can be expressed as 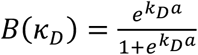. The Debye-Hückel electrostatic interactions of an ion pair act over a length scale of the order of *κ*^−1^, which is called the Debye screening length. *k*_*D*_ is related to ionic strength as,

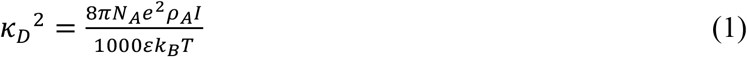

where *N*_*A*_ is the Avogadro number, *e* is the charge of an electron, *ρ*_*A*_ is the solvent density, *I* denotes solvent ionic strength, *k*_*B*_ is the Boltzmann constant, and T is the temperature. To avoid overlap among beads, a steep repulsion interaction has been defined as 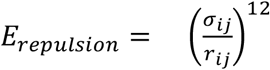 with *σ*_*ij*_=4 Å. Hydrophobic PHE residues interact through a short-range 12-10 dispersion interaction

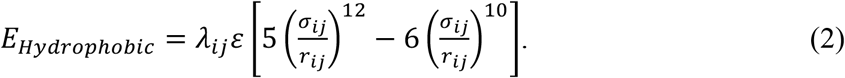

Phe–Phe interactions were described by a short-range potential with ε = 0.2 kcal mol^−1^, that reproduces experimentally observed dimensions of intrinsically disordered peptides^42^. Langevin dynamics simulations were performed for 100 binary polymers over a range of mixing fractions and subcritical temperatures at an ionic strength of 0.04 M using an implicit solvent. Two independent trajectories of 10^7^ integration steps were generated for each condition, and equilibrated largest condensates identified through cluster analysis were used for all structural, thermodynamic, and dynamical analyses.

### Stability of condensates, Criticality and Structural Morphology

To determine how differential sequence hydrophobicity regulates condensate stability, we constructed phase diagrams for heterotypic peptide mixtures across a range of mixing fractions (*X*_*High*−*ϕ*_) and compared them with homotypic condensates of identical average hydrophobicity (ϕ = 0.45) (Figure 1A–D, convergence of densities from independent runs has been shown in Figure S1). The largest equilibrium condensate was identified by cluster analysis, and critical temperatures were extracted from coexistence densities using universal critical scaling, enabling quantitative assessment of sequence-dependent phase stability. Critical temperature of condensates has been evaluated by

**Figure 1.**
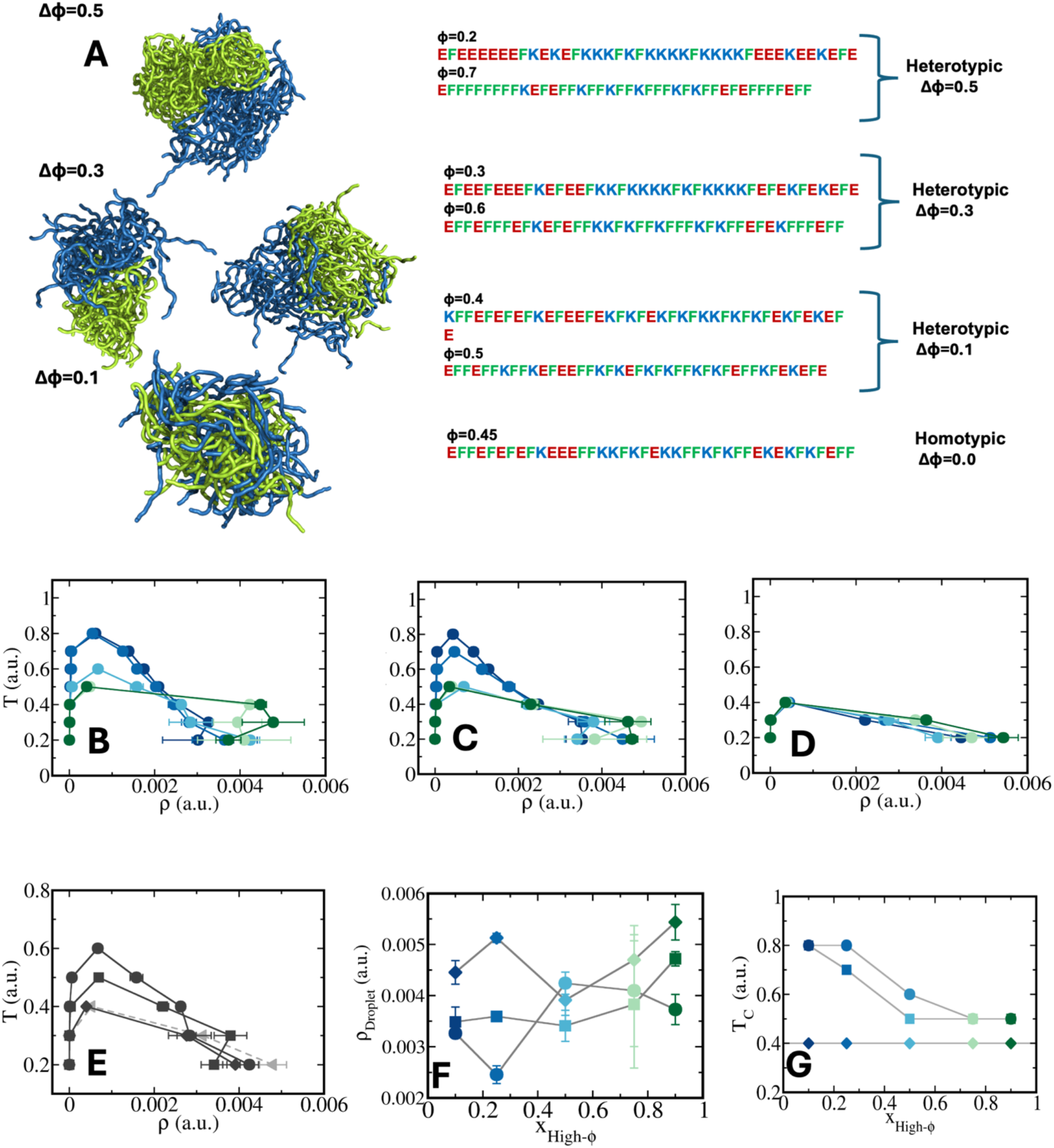
(A) Engineered condensates from designed sequence pairs with differential hydrophobicity enrichment. Blue and green colours represent peptides with dominant electrostatic and hydrophobic interactions respectively. Heterotypic pairs with hydrophobicity fraction ϕ = 0.2 and 0.7 (Δϕ = 0.5), ϕ = 0.3 and 0.6 (Δϕ = 0.3), and ϕ = 0.4 and 0.5 (Δϕ = 0.1) has been shown alongside the homotypic sequence with ϕ = 0.45 (Δϕ = 0.0), Coexistence curves have been shown for peptide pairs with differential hydrophobicity (B) Δϕ = 0.5, (C) Δϕ = 0.3, and (D) Δϕ = 0.1, respectively at different mixing fractions. (E) Phase diagrams for heterotypic condensate at *X*_*High*−*ϕ*_ =0.5 for all Δϕ variants compared to homotypic ones. (F) Droplet density as a function of mixing fraction *X*_*High*−*ϕ*_ at T/Tc =0.4 for all Δϕ variants. (G) Critical temperature (T_C_) as a function of mixing fraction (*X*_*High*−*ϕ*_). Colour codes represent mixing fractions (*X*_*High*−*ϕ*_). Circles, squares, diamonds and triangles represent Δϕ variants in descending order.

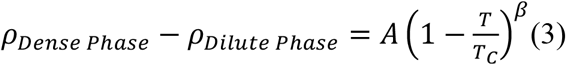

Where exponent β=0.3 for Ising model in three dimensions^63^.

Figure 1B–D reveals that condensate stability is strongly governed by sequence hydrophobicity mismatch and composition. For Δϕ = 0.5 and 0.3, critical temperatures decrease nearly twofold with increasing enrichment of the highly hydrophobic sequence (*X*_*High*−*ϕ*_), whereas condensates enriched in the less hydrophobic component remain substantially more stable. In contrast, for Δϕ = 0.1, the critical temperature is nearly insensitive to composition, reflecting minimal sequence asymmetry. The reduction in phase stability at high *X*_*High*−*ϕ*_, arises from a crossover from long-range electrostatic stabilization to predominantly short-range hydrophobic interactions. Notably, despite their lower critical temperatures, hydrophobicity-rich condensates exhibit markedly higher dense-phase packing below criticality.

To compare phase stability of heterotypic condensates to that of the homotypic ones we have shown phase diagrams for all three Δϕ peptide variants at *X*_*High*−*ϕ*_ = 0.50 with uniform homotypic condensate having sequence ϕ=0.45. Heterotypic condensates have nearly two-fold higher stability at extreme hydrophobicity mismatch than homotypic ones having same average sequence hydrophobicity of peptide pairs (Figure 1E and 1A). In addition we have observed density of condensates increases along enhancement of hydrophobicity rich sequences at the same distance from criticality T/T_C_=0.4 (Figure 1F). While peptide pairs having higher hydrophobicity mismatch form condensates with lower density, peptide pairs with minimal hydrophobicity mismatch has almost 1.5-2 times higher condensate density that too enhances with population of highly hydrophobic sequences in the system denoting efficient mixing in the droplet phase (Figure 1F). Variation of critical temperature has been shown for all the hydrophobicity mismatch sequence pairs at different mixing fractions (Figure 1G). Critical temperature drops gradually until *X*_*High*−*ϕ*_ = 0.25 for Δϕ=0.50 variant and thereafter, the decrease is rapid until *X*_*High*−*ϕ*_ = 0.75 as long-range electrostatic interactions get replaced by short-range hydrophobic ones due to enrichment of hydrophobic sequences in the system. T_C_ remains unchanged thereafter as short-range hydrophobic interactions dominate condensate formation. For Δϕ=0.30, the decrease in T_C_ is rapid until *X*_*High*−*ϕ*_ = 0.50 and then approaches constant upon increase in *X*_*High*−*ϕ*_ Sequence pair with Δϕ=0.10 showcase no effect of mixing fraction of peptides on criticality and shows exactly same T_C_ as that of ϕ=0.45 homogeneous condensate.

#### Energetics of condensation

The molecular basis of condensate stability was resolved by decomposing dense-phase energetics into hydrophobic and electrostatic contributions across sequence hydrophobicity mismatch (Δϕ), composition (*X*_*High*−*ϕ*_), and reduced temperature (Figure 2 and S2), revealing a sequence-dependent interplay between short- and long-range interactions. In addition, we have shown dissection into self (P1-P1 and P1-P2) and cross interactions (P1-P2) among peptides in Figure S3.

**Figure 2.**
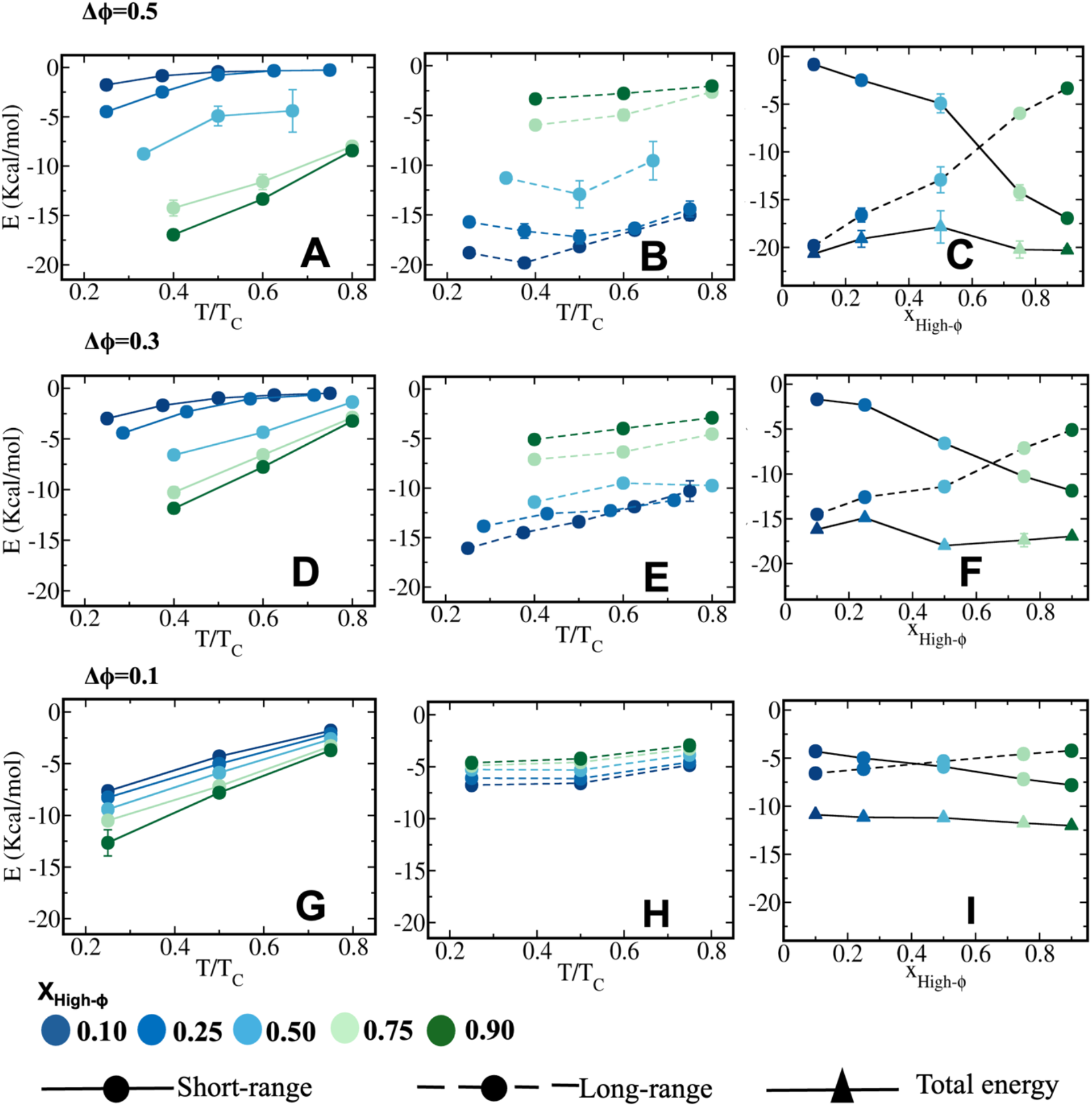
Long-range electrostatic and short-range hydrophobic energy contributions in stabilizing heterotypic condensates for (A-C) Δϕ = 0.5, (D-F) Δϕ = 0.3 and (G-I) Δϕ = 0.1. Colour codes represent mixing fractions (*X*_*High*−_*ϕ*). Panels A and B; D and E and G and H represent short and long-range contributions to energy of the droplet phase along temperature for Δϕ = 0.5; 0.3 and 0.1 variants respectively. Panels C, F and I showcase comparative contributions of electrostatic and hydrophobic interactions in stabilizing droplets at constant distance from criticality T/T_C_=0.4 along *X*_*High*−_*ϕ* for Δϕ = 0.5, 0.3 and 0.1 variants respectively; total energy of the droplet has been shown with triangles.

For the largest hydrophobic mismatch (Δϕ = 0.5; Figure 2A–C), dense-phase stabilization is dominated by short-range hydrophobic interactions at higher *X*_*High*−*ϕ*_. Hydrophobic stabilization is maximal (~17 kcal mol^−1^) in hydrophobicity-rich condensates (*X*_*High*−*ϕ*_ = 0.9) at low temperature (T/T_C_ = 0.2) and decreases with both increasing temperature and decreasing *X*_*High*−*ϕ*_, reaching ~10 kcal mol^−1^ at equimolar composition and becoming negligible near the critical point. These results demonstrate that hydrophobic cohesion scales directly with the abundance of hydrophobicity-rich sequences.

Electrostatic stabilization displays the opposite trend (Figure 2B), reaching ~17 kcal mol^−1^ in low-hydrophobicity-rich condensates (*X*_*High*−*ϕ*_ = 0.1). Unlike hydrophobic interactions, electrostatic contributions remain substantial (~10–12 kcal mol^−1^) near the critical point, reflecting their long-range character and persistent intermolecular association. At T/Tc = 0.4, condensate stabilization undergoes a composition-driven crossover from electrostatic to hydrophobic dominance, with the transition occurring near *X*_*High*−*ϕ*_ = 0.5 (Figure 2C). A similar but attenuated trend is observed for intermediate sequence heterogeneity (Δϕ = 0.3; Figure 2D–F). Hydrophobic stabilization decreases with increasing temperature and decreasing *X*_*High*−*ϕ*_, whereas electrostatic stabilization remains comparatively robust and weakly temperature dependent. These results highlight the greater thermal sensitivity of hydrophobic interactions relative to long-range electrostatic interactions. At T/T_C_ = 0.4, the dominant driving force switches from electrostatic to hydrophobic interactions near *X*_*High*−*ϕ*_ = 0.25, marking a composition-dependent crossover in condensate stabilization (Figure 2F). For minimal sequence heterogeneity (Δϕ = 0.1; Figure 2G–I), hydrophobic and electrostatic interactions contribute comparably to condensate stabilization. Their complementary presence largely compensate each other, yielding nearly composition-independent stabilization. This energetic compensation promotes peptide miscibility and suppresses interaction-driven segregation (Figure 2I).

### Heterogeneous Translational Diffusion in Droplet Phase

To elucidate dynamics in multi-phasic condensates formed by the heterotypic sequence pairs with hydrophobicity mismatch we computed mean squared displacements (MSD) of peptides in condensate phase as follows:

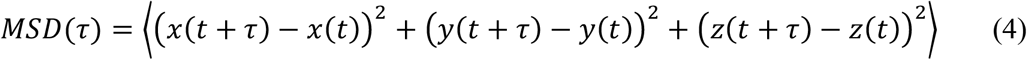

Mean-squared displacement analysis at T/Tc = 0.4 reveals a pronounced composition-dependent dynamical asymmetry for peptide pairs with large hydrophobicity mismatch (Δϕ = 0.3–0.5; Figure S4 (A-F). At low *X*_*High*−*ϕ*_, low-ϕ peptides are less mobile than high-ϕ counterparts, whereas their mobilities reverse in hydrophobicity-rich condensates. This dynamical inversion originates from sequence-dependent spatial segregation, which generates distinct local environments and heterogeneous molecular mobility.

At significantly lower hydrophobicity mismatch (Δϕ=0.1), we observed high-ϕ peptides irrespective of varying mixing fraction has lower mean-squared displacement than Low-ϕ ones in every scenario (Figure S4 (G-I)). Convergence of representative MSD profiles has been shown from independent runs in Figure S5.

To quantify condensate fluidity, we analysed the diffusivity ratio 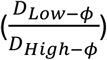, as a function of reduced temperature (Figure 3). Diffusivity is related to MSD as *MSD*(*τ*) = 6*Dτ*. The diffusivity ratio increases systematically with enhancement of *X*_*High*−*ϕ*_, revealing a composition-driven inversion of peptide mobility. For large sequence heterogeneity (Δϕ = 0.3– 0.5) and low *X*_*High*−*ϕ*_, high-ϕ peptides diffuse faster in electrostatically stabilized condensates, whereas low-ϕ peptides become up to threefold more mobile at high *X*_*High*−*ϕ*_ regime. Such mobility crossover indicates a reorganization of condensate architecture into distinct core–shell morphologies, consistent with the energetic and structural analyses (Figure 3A-B and Figure 2A-F and Figure 3G).

**Figure 3.**
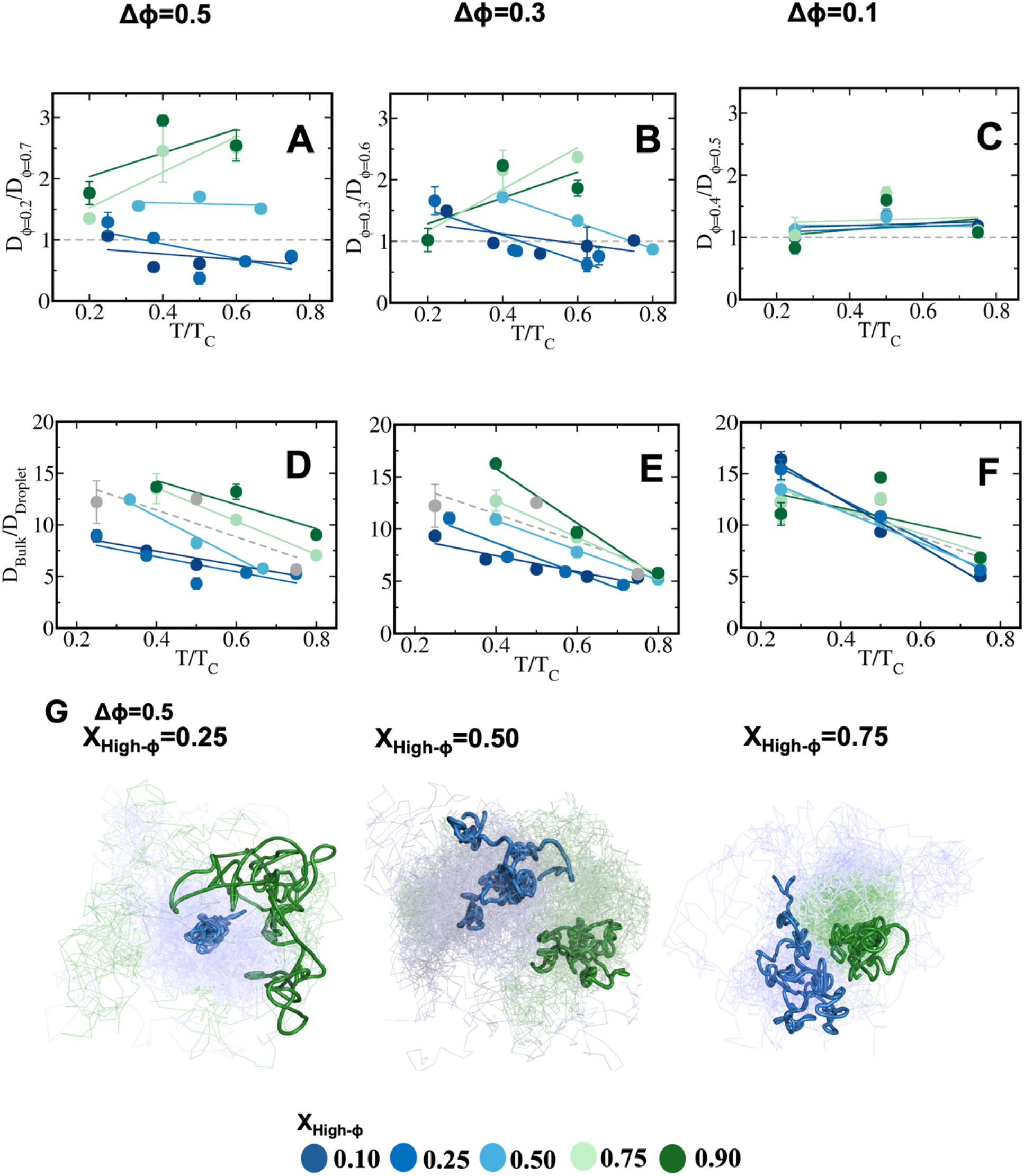
Differential dynamics in the condensate phase. Ratio of diffusivity among low and high-ϕ peptides in the dense phase 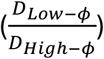 as a function of temperature for designed systems having (A) Δϕ = 0.5, (B) Δϕ = 0.3, and (C) Δϕ = 0.1 at different mixing fractions (*X*_*High*−*ϕ*_). Peptide diffusivity in bulk relative to condensate phase 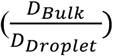 as a function of T/T_C_ for systems with (D) Δϕ = 0.5, (E) Δϕ = 0.3, and (F) Δϕ = 0.1 at various *X*_*High*−*ϕ*_. Colour codes represent mixing fractions (*X*_*High*−*ϕ*_). Grey represent corresponding homotypic reference (ϕ = 0.45). (G) Representative visual trajectories of peptides in condensate phase at significantly high hydrophobicity mismatch Δϕ = 0.5 and at different mixing fractions (*X*_*High*−*ϕ*_). Blue and green colours represent peptide trajectories with dominant electrostatic and hydrophobic interactions respectively.

At relatively higher *X*_*High*−*ϕ*_, the core of the droplet is formed by peptides with dominant hydrophobic interactions and low-ϕ peptides diffuse on the surface of the droplet as also evident from energetic analysis of condensation, resulting in 2-3 fold enhanced diffusivity of the low-ϕ ones compared to high-ϕ ones (Figure 3A and B and Figure 2A-B, D-E and Figure 3G).

At relatively low heterogeneity of the peptide pairs (Δϕ=0.1), low and high-ϕ peptides form a nearly homogeneous droplet having uniform diffusivity (Figure 3C).

To demonstrate liquid-like nature of the droplets formed, we quantified the ratio of bulk peptide diffusivity to that of droplets (Figure 3D-F). 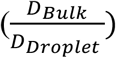 showcases prominent enhancement upon increase in mixing fraction at T/T_C_=0.4 for highly heterotypic sequence pairs namely Δϕ=0.3-0.5 (Figure 3D-E). Bulk diffusivity is nearly 5-7 times higher than condensates at relatively low mixing fraction (*X*_*High*−*ϕ*_). At relatively higher mixing fraction (*X*_*High*−*ϕ*_), condensate dynamics drops and ranges up to 17 folds slower than that of bulk peptide diffusivity. 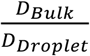 diminishes linearly as condensates approach criticality. While at low *X*_*High*−*ϕ*_ limit, bulk diffusivity is only 5-times higher than dense phase near criticality (T/T_C_=0.7-0.8), at high *X*_*High*−*ϕ*_ limit the same is 5-10 folds faster than droplet depending upon Δϕ. 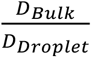 in homotypic condensate varies 8-14 folds as function temperature scaled to criticality.

At even lower heterogeneity of the sequence pairs, the liquid-like nature is fairly similar for all the mixing fractions i.e. ranging 12-15 times slower than bulk far away from criticality and as a function of mixing fraction (Figure 3F).

Representative visuals showcase trajectories of selected peptides in heterotypic multiphase condensates (Figure 3G) for Δϕ=0.5 at *X*_*High*−*ϕ*_=0.25, 0.50 and 0.75 at T/T_C_=0.5. Within a single condensate, multiple spatial regions enriched with trajectories of either High or low-ϕ peptides are prominent signifying differential liquid-like nature within one large condensate (Figure 3G). While trajectories of high-ϕ and low-ϕ peptides are well mixed in Δϕ=0.1 condensates, the same is distinctly demixed in Δϕ=0.5 at T/T_C_=0.5.

In order to understand the correlation between translational dynamics and single molecule conformational motions of peptides having differential hydrophobicity in condensate phase, we computed time-correlation function of end-to-end distance for peptide-pairs and plotted the average timescale of relaxation for designed peptide pairs as a ratio at different mixing fractions. Time correlation function has been defined as

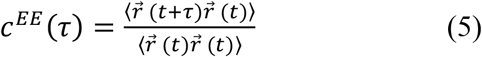

where 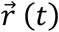 represents end-to-end distance vector of peptides (Figure S6). Convergence of time correlation functions has been shown from independent runs in Figure S7.

Single-chain conformational dynamics mirror the translational mobility (Figure 4A-C and Figure 3A-C). For Δϕ = 0.5, high-ϕ peptides reconfigure nearly twofold faster than low-ϕ peptides at low *X*_*High*−*ϕ*_, whereas increasing *X*_*High*−*ϕ*_, reverses this trend, with high-ϕ peptides becoming 2–3-fold slower. This inversion reflects a composition-driven reorganization of condensate architecture: electrostatically enriched condensates favour hydrophobic peptides at the interface, while hydrophobicity-rich condensates confine high-ϕ peptides within a densely packed core, restricting their conformational freedom (as observed in Figure 1A and 3G snapshots).

**Figure 4.**
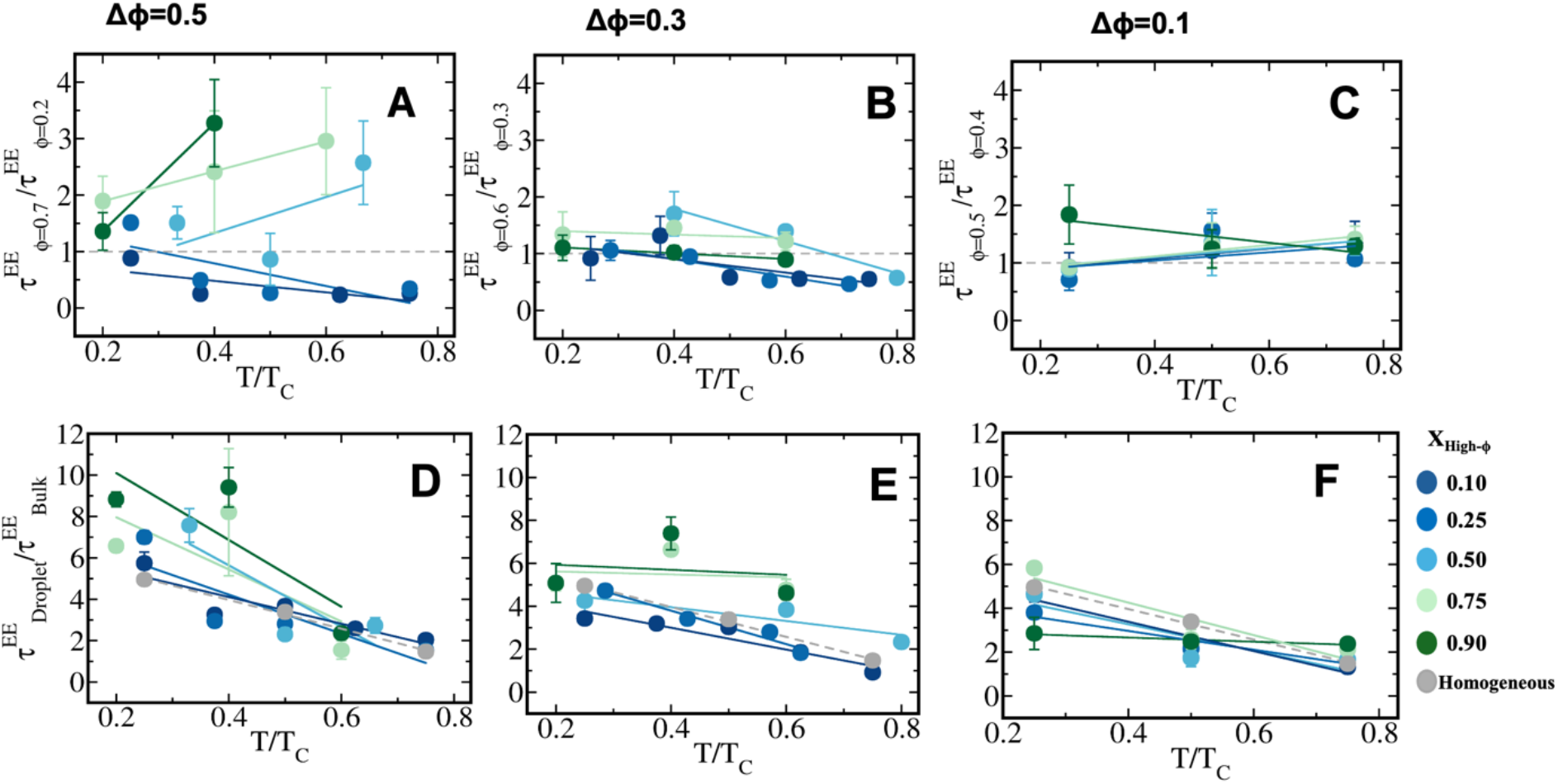
Chain reconfiguration dynamics in heterotypic condensates. Ratio of Reconfiguration lifetime 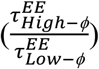 among low and high-ϕ peptides in condensate phase as a function of temperature for heterotypic systems with (A) Δϕ = 0.5, (B) Δϕ = 0.3, and (C) Δϕ = 0.1, respectively at various 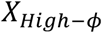. Average chain reconfiguration lifetime in the condensate phase compared to the bulk phase 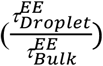 as a function of temperature for heterotypic systems with (D) Δϕ = 0.5, (E) Δϕ = 0.3, and (F) Δϕ = 0.1 respectively. Colour codes represent mixing fractions (*X*_*High*−*ϕ*_). Grey represent corresponding homotypic reference (ϕ = 0.45).

Reducing the hydrophobicity mismatch (Δϕ = 0.3, Figure 4B) attenuates the single molecule conformational dynamical asymmetry. Although low-ϕ peptides remain slower at low *X*_*High*−*ϕ*_, increased intercalation and comparable hydrophobic and electrostatic stabilization creates similar local environments, resulting in nearly indistinguishable conformational dynamics at higher hydrophobic-rich compositions (Figure 2D-F). This convergence is even more pronounced for Δϕ = 0.1, where extensive mixing eliminates sequence-dependent dynamical differences (Figure 4C, Figure 2G-I).

While comparing peptide dynamics in droplet and dilute bulk, we observed peptide conformational motions slow down heavily with the enrichment of hydrophobic peptides. At significant mismatch of peptide hydrophobicity (Δϕ=0.5, Figure 4D), we have observed peptides conformal dynamics in dense phase is nearly 4-6 folds slower than bulk depending upon temperature at far away from criticality at low *X*_*High*−*ϕ*_. At enhanced mixing fraction of hydrophobic peptides, conformational slowdown ranges up to 6-10 times far away from criticality (T/T_C_=0.3). At moderate mismatch of peptide hydrophobicity (Δϕ=0.1-0.3, Figure 4E-F), conformational dynamics of dense phase peptides is only 4-6 folds slower at significant enrichment of hydrophobic peptides while the same is only 2-4 folds slower in condensates having enrichment of peptides with stronger electrostatics near T/T_C_=0.3. Conformational dynamics of peptides in homogeneous droplet has been shown for reference with grey spheres showing 4-5 folds slowdown w.r.t bulk in resemblance with peptide pairs with lower heterogeneity.

At constant relative temperature (T/T_C_=0.4), bulk diffusion is 8 fold faster than dense-phase diffusion for Δϕ=0.5 at low *X*_*High*−*ϕ*_, increasing to 13-16 times with hydrophobic enrichment. Consequently, 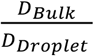 increases modestly up to *X*_*High*−*ϕ*_ = 0.25, where electrostatics dominate, and rises sharply thereafter as hydrophobic interactions increasingly arrest condensate dynamics. In contrast, for Δϕ =0.1-0.3, condensates remain relatively mobile up to *X*_*High*−*ϕ*_ = 0.50, followed by a rapid increase in 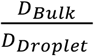 at higher compositions (Figure 5A). Consistently, 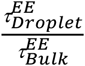 increases from 2-3 times to 6-9 times. Conformational relaxation exhibits a corresponding dynamical crossover for Δϕ =0.5, with fast peptide motions below *X*_*High*−*ϕ*_ = 0.25 and slow dynamics above this threshold. The crossover weakens for Δϕ =0.3 and disappears for Δϕ =0.1(Figure 5B).

**Figure 5.**
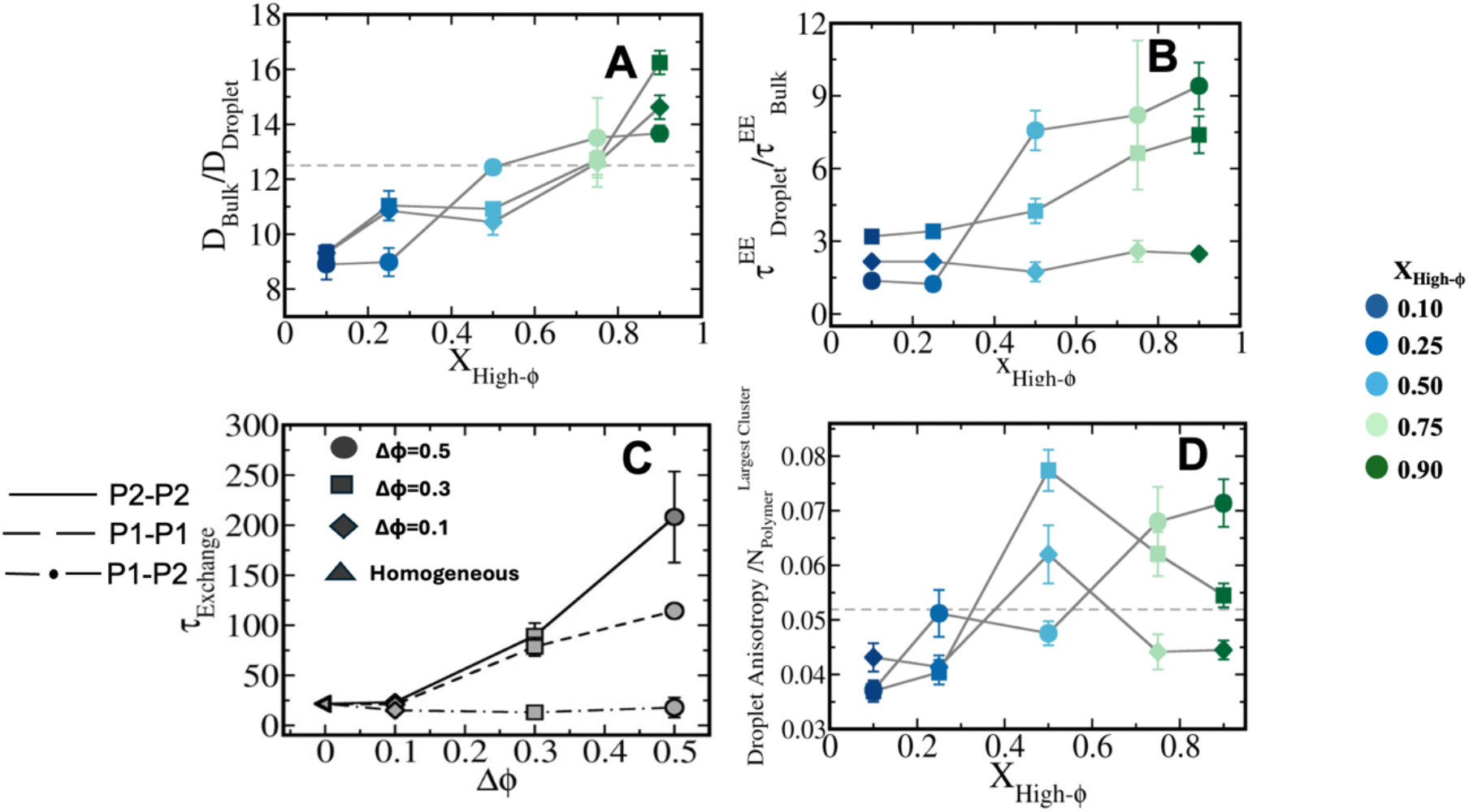
Composition-dependent exchange dynamics and condensate properties of heterotypic systems at (T/T_C_) = 0.4. (A) Peptide diffusivity in bulk compared to the condensate 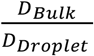 shown as a function of fraction of the high-ϕ polymers (*X*_*High*−*ϕ*_) at T/T_C_=0.4. (B) Chain reconfiguration lifetime in the condensate compared to bulk 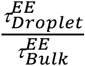 as a function of *X*_*High*−*ϕ*_ at T/T_C_=0.4. (C) Average neighbour exchange timescale (*τ*_Exchange_) for peptides as a function of the composition difference (Δϕ) for heterotypic systems at *X*_*High*−*ϕ*_ = 0.50 along sequence heterogeneity of binary peptide pairs (Δϕ). (D) Droplet anisotropy scaled to cluster size as a function of *X*_*High*−*ϕ*_ for all Δϕ variants. Colour codes represent mixing fractions. Circles, squares, diamonds and triangles represent Δϕ variants in descending order.

To correlate single-molecule conformational dynamics with neighbour exchange kinetics, we introduce a sticker exchange time correlation function

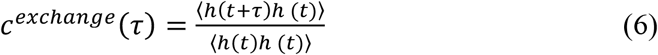

h(t) denotes a step function; h(t)=1 when two peptides are neighboring ones for a time frame and 0 otherwise. Time correlation functions have been plotted for peptide1-peptide1 (P1-P1, low-ϕ peptides), peptide2-peptide2 (P2-P2, high-ϕ peptides) and peptide1 and peptide2 (P1-P2, cross sticker interactions among low and high-ϕ peptides) (Figure S8). We have shown the average timescale of neighbor exchange kinetics at *X*_*High*−*ϕ*_ = 0.50 for all Δϕ variants along with the homogeneous one (Δϕ =0) (Figure 5C). We observed that P1-P1 (low-ϕ peptides) and P2-P2 (high-ϕ peptides) contacts have longer lifetime along enhancement of Δϕ indicating dominance of short-range hydrophobic and long-range electrostatic contacts at different spatio-temporal regions of droplets indicating multiphasic behavior confirmed by P1-P2 (cross sticker interactions among low and high-ϕ peptides) cross contact lifetime being negligible in each scenario (Figure 5C).

To correlate shape anisotropy of dense phase to internal morphology and dynamics, droplet shape anisotropy (p) has been defined as:

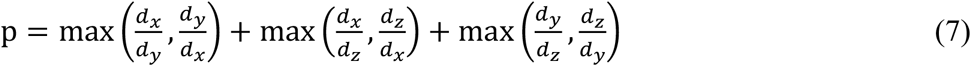

where *d*_*x*_, *d*_*y*_, *d*_*z*_ are the largest diameter in each dimension of the largest cluster. An ideal sphere has a value of 3 along this definition and deviation from the same indicates an anisotropic condensate. Phase separated condensates tend to acquire a spherical shape to reduce surface tension. Deviation of shape of a condensate from a spherical one may originate from competitive interactions. Droplet anisotropy increases with *X*_*High*−*ϕ*_, signalling multiphasic, less compact condensates at Δϕ=0.5. Below *X*_*High*−*ϕ*_ = 0.5, anisotropy remains lower than in the homogeneous condensate, consistent with a single-phase droplet and surface localization of hydrophobic peptides. For Δϕ=0.1-0.3, anisotropy peaks at *X*_*High*−*ϕ*_ = 0.5 owing to maximal multiphase formation and declines thereafter because of enhanced mixing and multiple smaller condensates (Figure 5D and S9).

## Conclusion

In summary, we demonstrate that sequence hydrophobicity mismatch alone constitutes a robust molecular design principle for programming the phase behaviour of heterotypic biomolecular condensates in absence of cation-π cross competing interactions. By systematically varying the hydrophobic disparity and composition of engineered peptide pairs, we uncover how the competition between short-range hydrophobic interactions and long-range electrostatic forces simultaneously governs condensate thermodynamic stability, internal morphology, and molecular dynamics. Beyond modulating phase stability, sequence heterogeneity gives rise to emergent multiphasic organization, where composition-dependent inversion of core–shell architecture produces spatially distinct interaction environments within a single condensate. Remarkably, heterotypic condensates are consistently more stable than homotypic condensates possessing the same average sequence hydrophobicity, highlighting the cooperative and non-additive nature of intermolecular interactions in multicomponent systems. This structural organization is directly reflected in the transport and conformational dynamics of individual peptide species, establishing a clear link between sequence chemistry, mesoscale compartmentalization, and condensate material properties.

These results provide a unified molecular picture connecting sequence-encoded interactions to the thermodynamics and dynamics of heterogeneous condensates while offering predictive design rules for engineering synthetic biomolecular assemblies. More broadly, our findings suggest that tuning interaction heterogeneity, rather than simply the average interaction strength, represents a powerful strategy for controlling condensate architecture and function. We anticipate that these principles will facilitate the rational design of programmable peptide condensates with tailored compartmentalization, transport characteristics, and mechanical properties for applications in synthetic biology, biomolecular engineering, and responsive soft materials.

### Supplementary data

Convergence of droplet density from independent runs (S1), Total energetic stability of droplet phase along temperature for all mixing fractions for all Δϕ variants (S2), Self and cross energetic components among peptides along temperature and Δϕ variants (S3), Average mean squared displacements (MSD) of low and high-ϕ peptides at chosen *X*_*High*−*ϕ*_ at T/T_C_=0.4 for all Δϕ variants (S4) at *X*_*High*−*ϕ*_ = 0.25,0.50 *and* 0.75, Representative convergence of MSD profiles for all Δϕ variants at *X*_*High*−*ϕ*_ = 0.50 for high and low-ϕ peptides (S5). Average end-to-end distance time correlations of low and high-ϕ peptides at chosen *X*_*High*−*ϕ*_ at T/T_C_=0.4 for all Δϕ variants (S6). Representative convergence of end-to-end dynamics time correlation functions for all Δϕ variants at *X*_*High*−*ϕ*_ = 0.50 for high and low-ϕ peptides (S7), Neighbour exchange time correlation functions for self and cross peptide contacts (S8), Variation of droplet anisotropy along temperature for *X*_*High*−*ϕ*_ = 0.25,0.50,0.75 for all Δϕ variants (S9). Droplet enrichment and bulk to droplet energetic and translational entropic gain (S10).

## Notes

### Competing Interest Statement

The authors have declared no competing interest.

### Funding

- ANRF: ANRF/ECRG/2024/000734/LS
- Indian Institute of Technology Jodhpur: I/SEED/MKZ/20230169

