## Supplementary data for "Engineering Heterotypic Biomolecular Condensates with Synthetic Peptides for Controlled Spatial Organization and Liquid-like Nature"

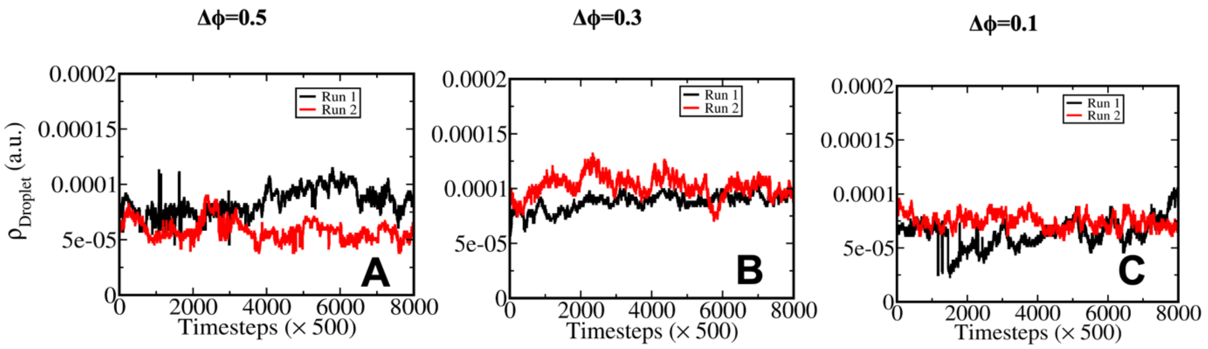


**Figure S1.** Convergence of droplet density obtained from two independent simulation runs at mixing composition $X_{\mathrm{High}-\phi} = 0.50$. Droplet density is shown as a function of simulation time for (A) $\Delta\phi = 0.5(\phi = 0.2$ and $0.7)$, (B) $\Delta\phi = 0.3(\phi = 0.3$ and $0.6)$, and (C) $\Delta\phi = 0.1(\phi = 0.4$ and $0.5)$, respectively. Black and red curves represent Run 1 and Run 2, respectively, demonstrating reproducibility and convergence of droplet density across independent simulations.

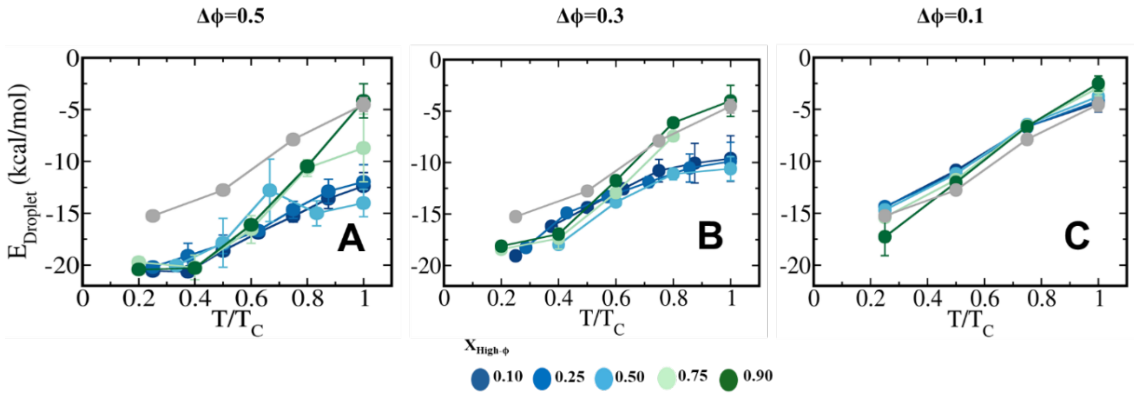


**Figure S2.** Average droplet interaction energy ($E_{\mathrm{Droplet}}$) as a function of temperature scaled by critical temperature for heterotypic condensates. Panels (A–C) correspond to the $\Delta\phi = 0.5(\phi = 0.2$ and $0.7)$, $\Delta\phi = 0.3(\phi = 0.3$ and $0.6)$, and $\Delta\phi = 0.1(\phi = 0.4$ and $0.5)$ systems, respectively. Different colours represent mixing fractions ranging from $X_{\mathrm{High}-\phi} = 0.10$ to 0.90, while grey symbols denote corresponding homotypic reference system (Figure 1A). Comparison illustrates how droplet interaction energy varies with temperature and composition for different degrees of sequence heterogeneity.

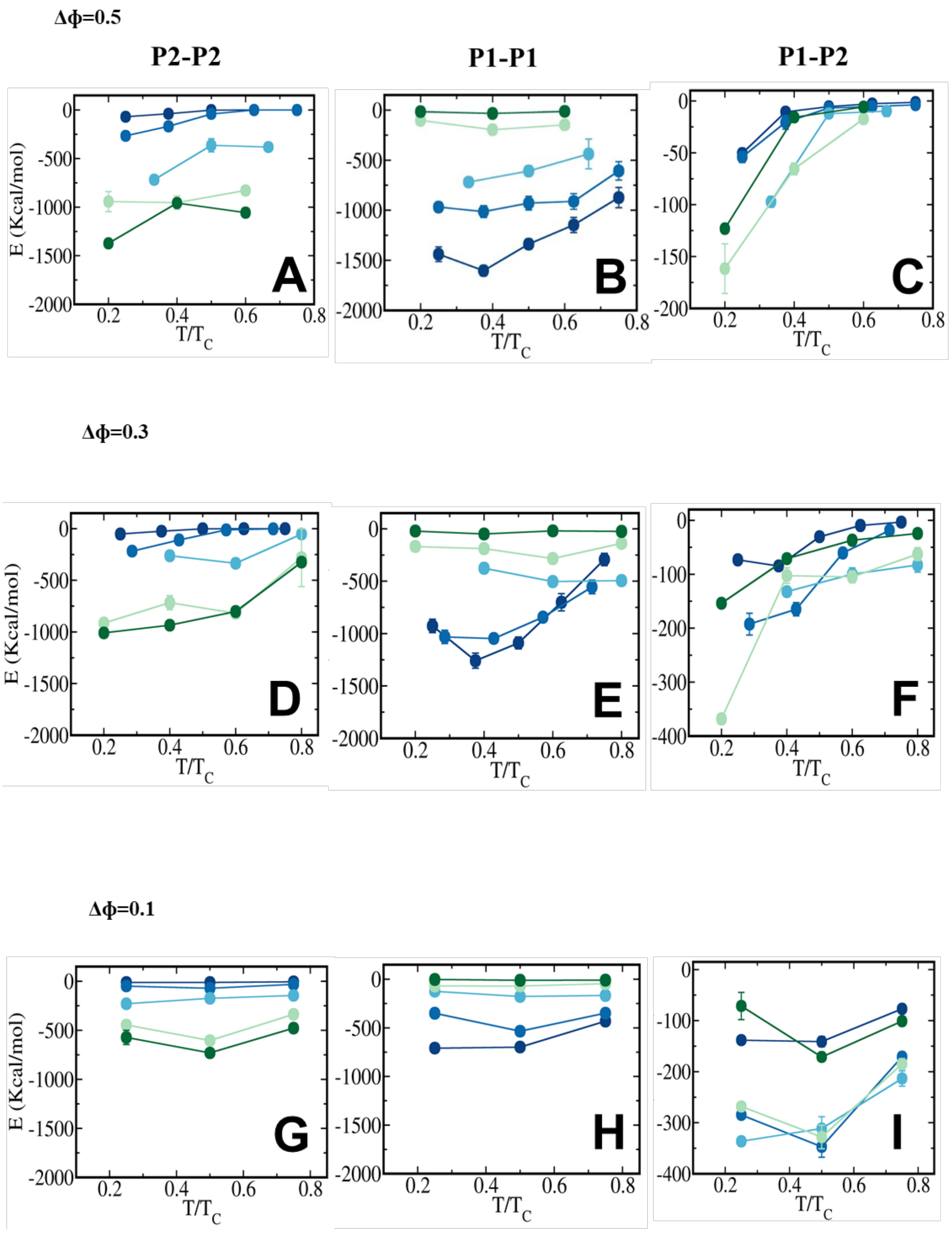


**Figure S3.** Dissection of total interaction energy into self- and cross peptide energy components in heterotypic condensates with (A–C) $\Delta\phi = 0.5$, (D–F) $\Delta\phi = 0.3$, and (G–I) $\Delta\phi = 0.1$. Panels (A), (D), and (G) show average self-interaction energy among high-$\phi$ (P2–P2) peptides experienced by each peptide in condensate phase for descending $\Delta\phi$ respectively. Panels (B), (E), and (H) show average self-interaction energy among low-$\phi$ (P1–P1) peptides. Panels (C), (F), and (I) show average cross-interaction energy between high-$\phi$ and low-$\phi$ (P1–

P2) peptides in condensate phase. Different high-$\phi$ compositions are shown in each panel, ranging from $X_{\text{High}-\phi}$ = 0.10 to 0.90, with colour changing from blue to green.

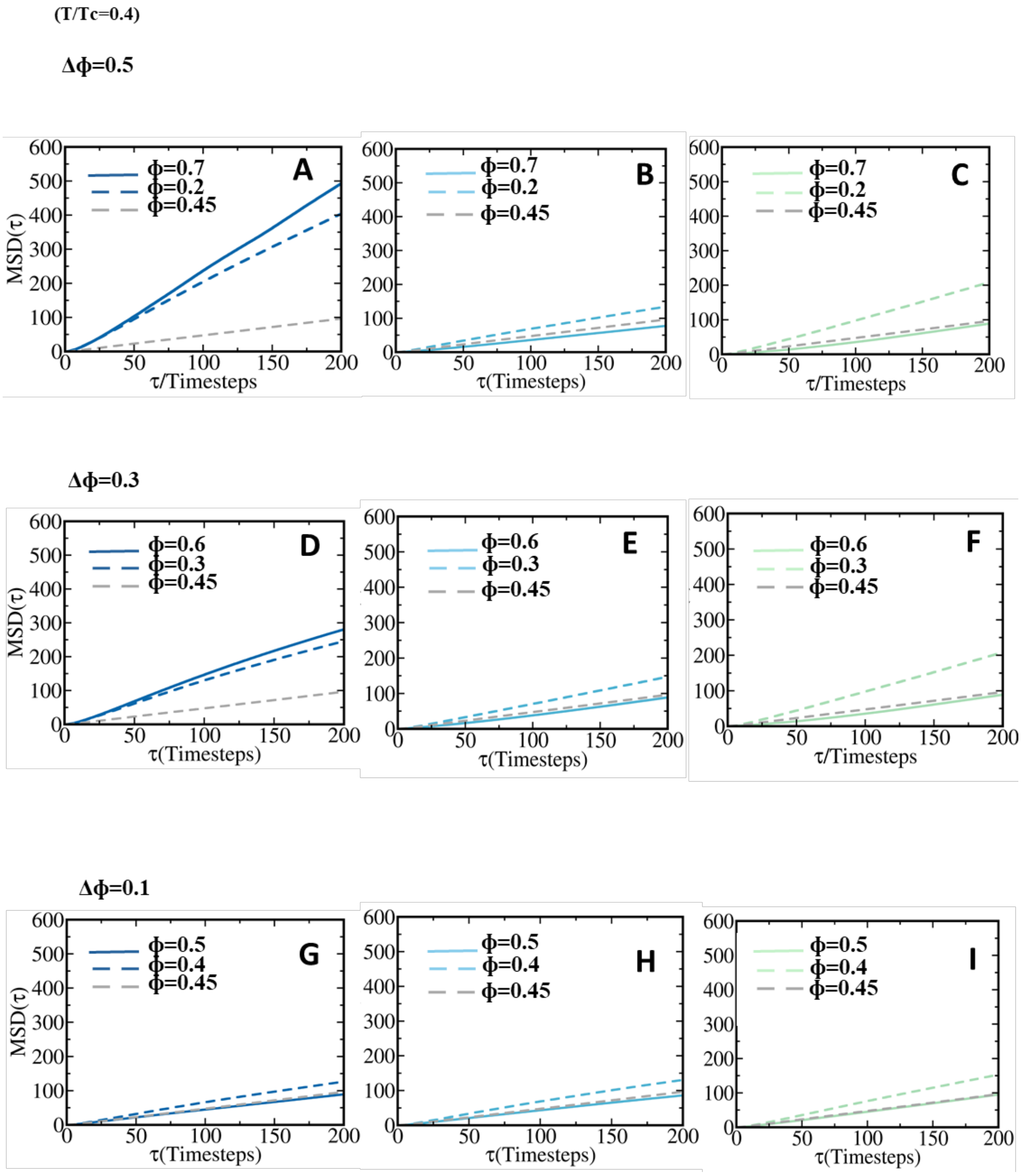


**Figure S4.** Mean squared displacement (MSD) of peptides in the condensate phase at $T/T_C$ = 0.4. Panels (A–C) correspond to the $\Delta\phi$ = 0.5 system (peptide $\phi$ = 0.2 and 0.7), panels (D–F) correspond to the $\Delta\phi$ = 0.3 system ($\phi$ = 0.3 and 0.6), and panels (G–I) correspond to the $\Delta\phi$ = 0.1system ($\phi$ = 0.4 and 0.5). Panels (A), (B), and (C) correspond to $X_{\text{High}-\phi}$ = 0.25, 0.50, and 0.75, respectively. Similarly, panels (D), (E), and (F), and panels (G), (H), and (I)

correspond to ($X_{\text{High}-\phi}$ = 0.25, 0.50, and 0.75), respectively. Solid lines represent the high-$\phi$ peptides, dashed lines represent the low-$\phi$ peptides, and the grey dashed lines denote the corresponding homotypic reference ($\phi$ = 0.45). The color scale corresponds to the peptide composition in each heterotypic system.

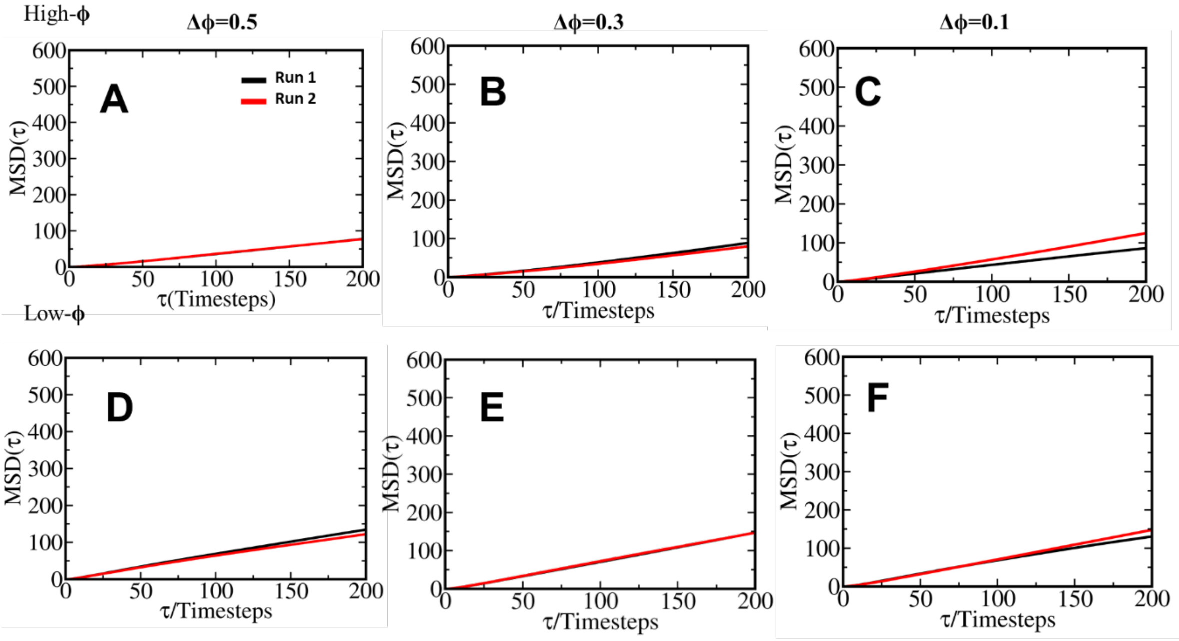


**Figure S5.** Convergence of mean squared displacements (MSD) of high-$\phi$ peptides obtained from two independent simulation runs at mixing composition $X_{\text{High}-\phi}$ = 0.50 and $T/T_C$=0.4 for (A) $\Delta\phi = 0.5(\phi = 0.2$ and 0.7), (B) $\Delta\phi = 0.3(\phi = 0.3$ and 0.6), and (C) $\Delta\phi = 0.1(\phi =$ 0.4 and 0.5), respectively. Panels (D–F) show corresponding MSD of low-$\phi$ peptides for designed systems with (D) $\Delta\phi = 0.5(\phi = 0.2$ and 0.7), (E) $\Delta\phi = 0.3(\phi = 0.3$ and 0.6), and (F) $\Delta\phi = 0.1(\phi = 0.4$ and 0.5), respectively. Black and red curves represent Run 1 and Run 2, respectively, demonstrating reproducibility and convergence of peptide dynamics across independent simulations.

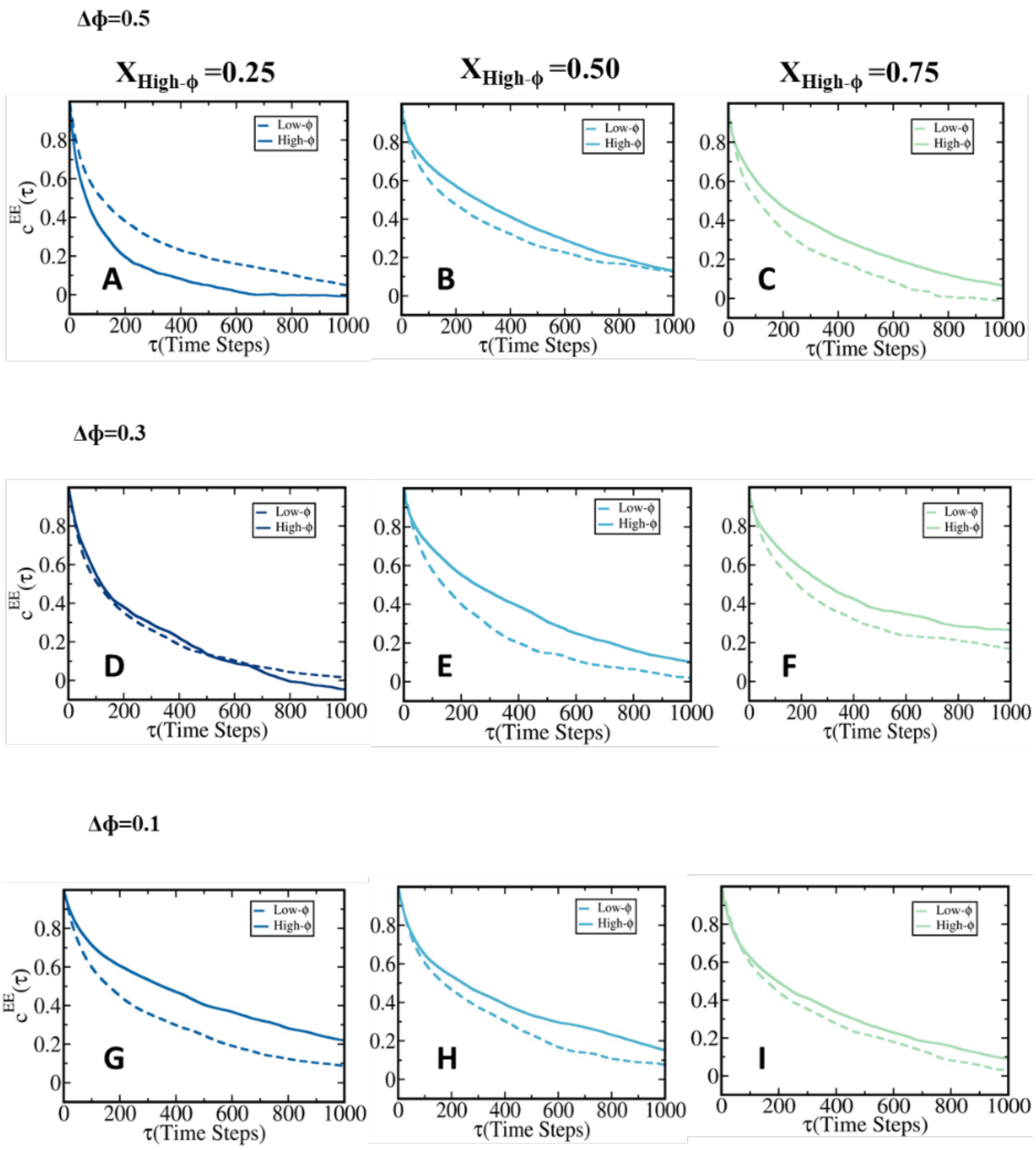


**Figure S6.** Representative end-to-end distance time correlation functions of high-$\phi$ and low-$\phi$ peptides in heterotypic condensates at different mixing compositions. Panels (A–C) correspond to the $\Delta\phi = 0.5$system ($\phi = 0.2$ and 0.7), panels (D–F) correspond to the $\Delta\phi = 0.3$system ($\phi = 0.3$ and 0.6), and panels (G–I) correspond to the $\Delta\phi = 0.1$system ($\phi = 0.4$ and 0.5). Within each row, panels from left to right correspond to mixing fraction $X_{\mathrm{High}-\phi} =$ 0.25, 0.50, and 0.75 respectively. Solid and dashed curves represent high-$\phi$ and low-$\phi$ peptides, respectively, illustrating the dependence of chain reconfiguration dynamics on sequence composition and mixing fraction.

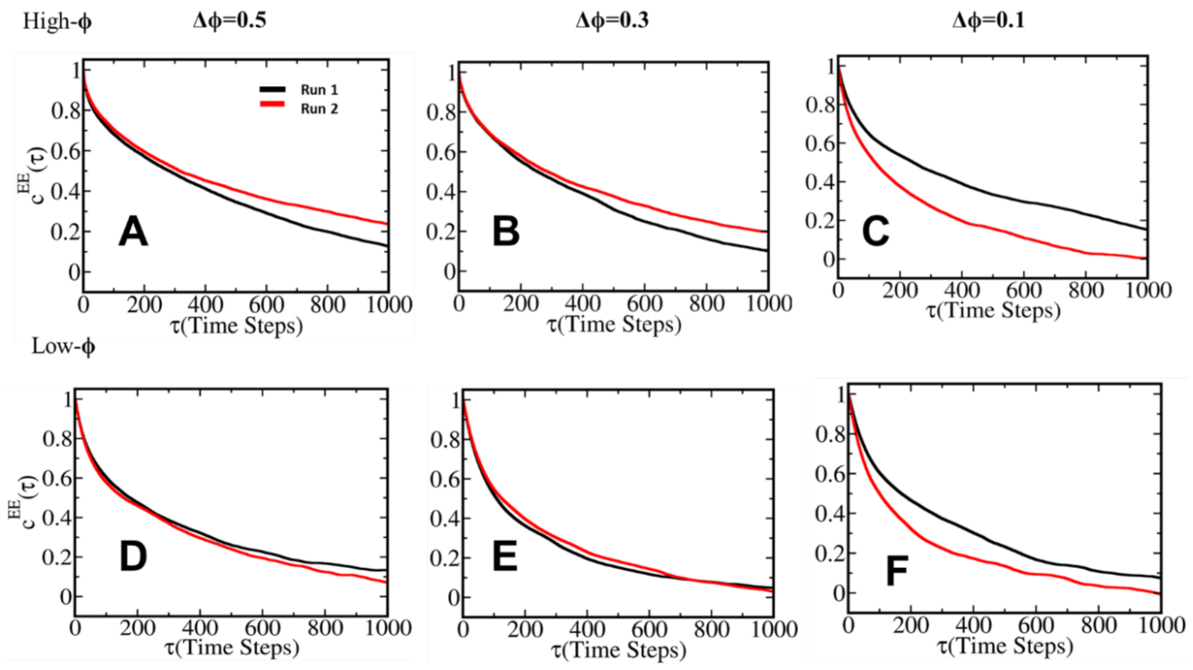


**Figure S7.** Representative end-to-end distance time correlation functions obtained from two independent simulation runs at mixing composition $X_{\text{High}-\phi} = 0.50$ and T/$T_C$=0.4. Time correlation functions have been shown for high-$\phi$ peptides for the designed systems (A)$\Delta\phi = 0.5(\phi = 0.2$ and 0.7), (B)$\Delta\phi = 0.3(\phi = 0.3$ and 0.6), and (C) $\Delta\phi = 0.1(\phi = 0.4$ and 0.5), respectively. Panels (D–F) show corresponding time correlation functions of low-$\phi$ peptides for the same systems respectively. Black and red curves represent Run 1 and Run 2, respectively, demonstrating reproducibility and convergence of chain reconfiguration dynamics across independent simulations.

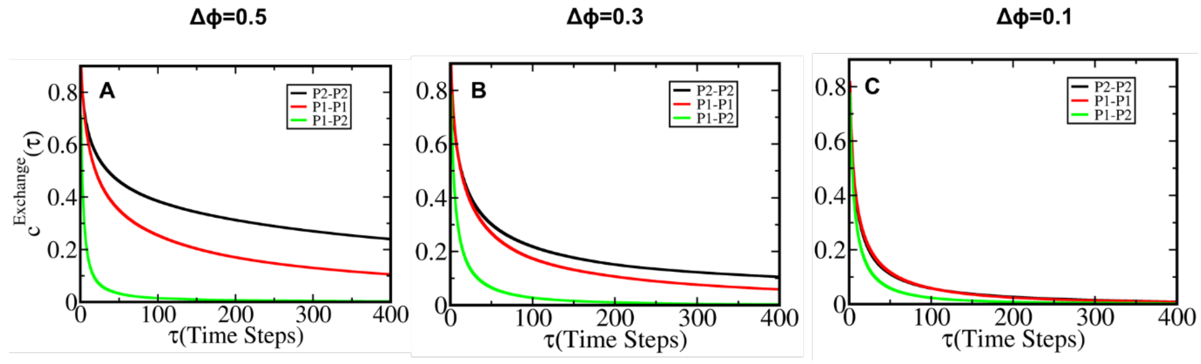


**Figure S8.** Representative neighbour contact exchange time correlation functions in condensate phase at $T/T_C = 0.4$ and $X_{\text{High}-\phi} = 0.50$. Panels (A–C) correspond to the $\Delta\phi = 0.5(\phi = 0.2$ and 0.7), $\Delta\phi = 0.3(\phi = 0.3$ and 0.6), and $\Delta\phi = 0.1(\phi = 0.4$ and 0.5) systems, respectively. In each panel, black, red, and green curves represent sticker exchange time correlation functions for high-$\phi$(P2–P2), low-$\phi$(P1–P1), low and high-$\phi$ (P1–P2) respectively.

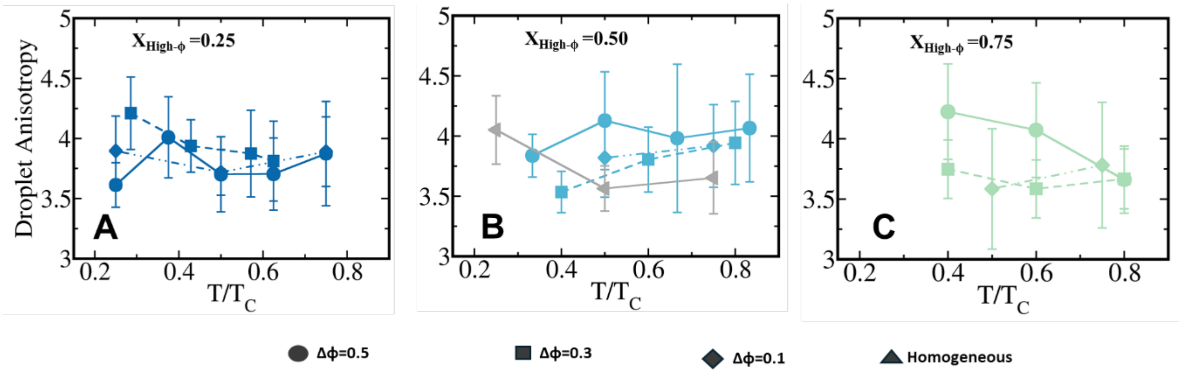


**Figure S9.** Droplet anisotropy as a function of temperature scaled to criticality for heterotypic condensates at fixed high-$\phi$ compositions: (A) $X_{\text{High}-\phi} = 0.25$, (B) $X_{\text{High}-\phi} = 0.50$, and (C) $X_{\text{High}-\phi} = 0.75$. In each panel, circles, squares, and diamonds represent heterotypic systems with $\Delta\phi = 0.5(\phi = 0.2$ and $0.7)$, $\Delta\phi = 0.3(\phi = 0.3$ and $0.6)$, and $\Delta\phi = 0.1(\phi = 0.4$ and $0.5)$, respectively. Grey symbols denote corresponding homotypic reference $(\phi = 0.45)$. Comparison illustrates effect of sequence heterogeneity on droplet shape anisotropy at identical high-$\phi$ compositions across three designed heterotypic systems.

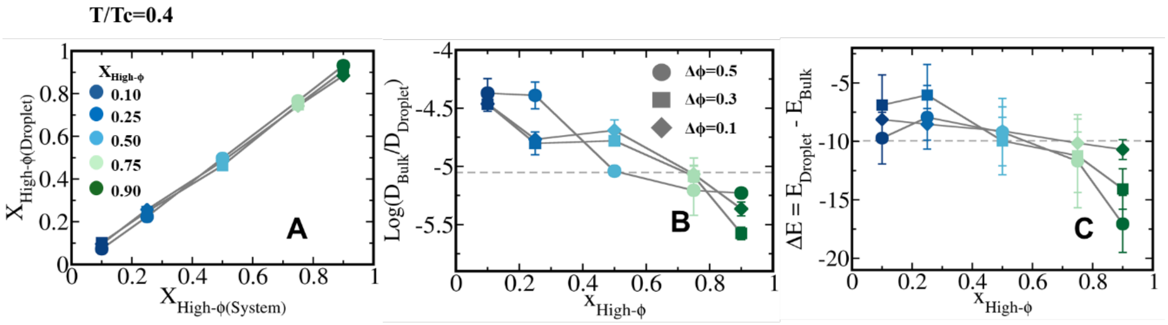


**Figure S10.** (A) Enrichment of high-$\phi$ peptides inside condensates $\left(X_{\text{High}-\phi}^{\text{Droplet}}\right)$ compared to overall system composition $\left(X_{\text{High}-\phi}^{\text{System}}\right)$ at $T/T_C = 0.4$ for heterotypic systems with $\Delta\phi = 0.5$, 0.3, and 0.1 respectively. (B) Log of bulk to condensate diffusivity ratio $(\log\frac{D_{Bulk}}{D_{Droplet}})$ , as a function of $X_{\text{High}-\phi}$ at $T/T_C = 0.4$ relating entropic advantage along Rosenfeld Scaling. Dashed horizontal line denotes reference homotypic scenario, $\log\frac{D_{Bulk}}{D_{Droplet}} = -5.05$. (C) Difference between average interaction energies in condensate and bulk phases, $\Delta E = E_{\text{Droplet}} - E_{\text{Bulk}}$, as a function of high-$\phi$ composition at $T/T_C = 0.4$. Dashed horizontal line indicates reference value for homotypic condensate, $\Delta E = -10$ kcal/mol. Circles, squares, and diamonds represent systems with $\Delta\phi = 0.5$, 0.3, and 0.1, respectively, while colour scale denotes high-$\phi$ composition ranging from $X_{\text{High}-\phi} = 0.10$ to 0.90.
